# Manipulation of carotenoid metabolism stimulates biomass and stress tolerance in tomato

**DOI:** 10.1101/2021.05.05.442770

**Authors:** Jose G. Vallarino, Jianing Mi, Ivan Petřík, Ondřej Novák, Sandra M. Correa, Monika Kosmacz, Michel Havaux, Manuel Rodriguez-Concepcion, Salim Al-Babili, Alisdair R. Fernie, Aleksandra Skirycz, Juan C. Moreno

## Abstract

Improving yield, nutritional value and tolerance to abiotic stress are major targets of current breeding and biotechnological approaches that aim at increasing crop production and ensuring food security. Metabolic engineering of carotenoids, the precursor of Vitamin-A and plant hormones that regulate plant growth and response to adverse growth conditions, has been mainly focusing on provitamin A biofortification or the production of high-value carotenoids. Here, we show that the introduction of a single gene of the carotenoid biosynthetic pathway in different tomato cultivars simultaneously improved photosynthetic capacity and tolerance to various abiotic stresses (e.g., high light, salt, and drought), caused an up to 77% fruit yield increase and enhanced fruit’s provitamin A content and shelf life. Our findings pave the way for developing a new generation of crops that combine high productivity and increased nutritional value with the capability to cope with climate change-related environmental challenges.

## INTRODUCTION

Climate change and the increasing world population are serious challenges facing world agriculture (Pareek et al., 2020). Indeed, current estimates indicate that food production should be doubled by 2050 (Ort et al., 2015; Xu, 2016). However, global warming and the anthropogenic activities that affect agricultural ecosystems and subsequent crop yield render this doubling a very difficult goal to achieve. Moreover, abiotic stresses, and especially salinity and drought, cause considerable crop losses, with yield reductions of almost 50% (Hussain et al., 2019; Roy et al., 2014). Therefore, a new generation of crops with enhanced fitness—as exemplified, for instance, by simultaneously improved photosynthetic efficiency, stress tolerance, and yield—are urgently needed to meet the desired levels of crop productivity. In the past decade, photosynthesis and photorespiration have been the preferred targets for manipulation to improve plant yield (Ding et al., 2016; Lopez-Calcagno et al., 2019; Simkin et al., 2017; Simkin et al., 2015; South et al., 2019; Timm et al., 2015). For example, two breakthrough genetic strategies for manipulating the xanthophyll cycle (manipulation of three genes) and glycolate metabolism (introduction of five genes) have documented increases in plant biomass of between 15% and 37%, respectively, in the cash crop tobacco (Kromdijk et al., 2016; South *et al*., 2019). However, to date, neither of these strategies have been demonstrated to work in food crops. Moreover, similar manipulation of the xanthophyll cycle in Arabidopsis resulted in a contradictory reduction in plant biomass (Garcia-Molina and Leister, 2020), bringing into question the general applicability of this method.

Another possibility for manipulating plant yield and fitness in crops might be provided by the carotenoids (e.g., β-carotene), which are isoprenoid pigments that rank among the most important plant secondary metabolites due to the diverse functions they fulfil in photosynthesis and signaling. Within chloroplasts, carotenoids like β-carotene and xanthophylls are key components of photosynthetic membranes and form pigment-protein complexes that are essential for photoprotection (Niyogi and Truong, 2013; Xu et al., 2020). β-carotene is also the precursor of abscisic acid (ABA) and strigolactones (SLs), so alterations in carotenoid content can affect hormone content and subsequent plant development and physiology (Al-Babili and Bouwmeester, 2015; Nambara and Marion-Poll, 2005). In recent years, new signaling and growth-promoting functions have been reported for carotenoid-derived molecules (commonly referred to as apocarotenoids), including β-cyclocitral (β-cc), dihydroactinidiolide (dhA), and zaxinone (Zax) (D’Alessandro et al., 2018; D’Alessandro et al., 2019; Dickinson et al., 2019; Hou et al., 2016; Wang et al., 2019). In animals, carotenoids consumed in the diet are also cleaved to produce retinoids (including vitamin A) and other molecules with signaling and health-promoting properties (Rodriguez-Concepcion et al., 2018). β-carotene is the main precursor of vitamin A in animals and the main precursor of several apocarotenoids and plant hormones in plants; therefore, increased accumulation of β-carotene might indirectly influence plant growth and development, as well as improve the nutritional value. β-carotene is produced by the action of lycopene β-cyclase (LCYB), indicating a potential for genetic manipulation of the expression of this gene as a two-for-one solution to improve both the fitness and the nutritional value of the chosen crop.

In our previous work, we expressed the LCYB-encoding *DcLCYB1* gene from carrot (*Daucus carota*) in tobacco and demonstrated growth-promoting and developmental effects of this gene (Moreno et al., 2020). Interestingly, these tobacco lines also showed enhanced tolerance to abiotic stresses, in addition to enhancement of biomass, yield, and photosynthetic efficiency (Moreno et al., 2021). These beneficial effects were mainly triggered by an enhanced accumulation of the phytohormones ABA and gibberellic acid (GA), but they were also a result of the greater photoprotection afforded by the accumulation of xanthophylls. We therefore hypothesized that any LCYB-encoding gene, independent of its origin (plant or bacterial), might trigger similar beneficial effects to those observed with the carrot *DcLCYB1* gene in tobacco (Moreno *et al*., 2020).

In the present study, we explored this hypothesis using previously generated tomato cultivars that overexpress three different *LCYB* genes (from plant and bacterial origins) following plastid or nuclear transformation. We confirmed that the overexpression of any *LCYB* gene is sufficient to trigger a molecular response that results in modulation of carotenoid (pro-vitamin A) and hormone content, with a subsequent alteration in plant architecture, photosynthetic efficiency, stress tolerance, and yield.

## RESULTS

### Tomato productivity under different environmental conditions

Given our recent findings that expression of the carrot *DcLCYB1* gene resulted in increased photosynthetic efficiency, photoprotection, stress tolerance, plant biomass, and yield in tobacco (Moreno *et al*., 2021; Moreno *et al*., 2020), we decided to evaluate whether manipulation of LCYB activity could confer similar growth advantages in an economically important food crop. We tested our hypothesis by exploiting the availability of several tomato cultivars overexpressing different LCYB-encoding genes. In particular, we used a Red Setter cultivar with a nuclear construct overexpressing a tomato LCYB (line H.C.) and two transplastomic lines expressing LCYB-encoding genes from daffodil in the IPA6+ background (line pNLyc#2) or from the bacterium *Erwinia uredovora* (renamed *Pantoea ananatis*) in the IPA6-background (line LCe) (see Materials and Methods; **Table S1**). Growth evaluation under different climate conditions (fully controlled, semi-controlled, and uncontrolled conditions) revealed robust and homogeneous changes in plant height (increased and reduced plant height for transplastomic and nucear lines, respectively) of the transgenic lines in comparison to their respective wild type in all climate conditions **(fig. S1)**. Due to the robustness of the phenotypes, we selected the semi-controlled conditions (greenhouse) to perform a detailed molecular and physiological characterization of this phenomenon. Interestingly, the pNLyc#2 and LCe transplastomic lines showed longer stems than their respective wild-type plants, thereby allowing a more spaced allocation of their leaves along the stem. By contrast, the H.C. nuclear line showed reduced plant height (**Fig. 1A-C**). In addition, leaves from pNLyc#2 were larger than the IPA6+ leaves, while leaves from the H.C. line were smaller than those from its wild type R.S. (**fig. S2A, D**). By contrast, leaves from the LCe line showed sizes similar to the wild type (**fig. S2G**). The fruit size was similar to the wild type in the pNLyc#2 line but was slightly larger in the LCe line (**fig. S2J**), while the fruit from the H.C. line were considerably larger when compared to those from its respective wild type (**fig. S2B, E, H, J)**.

**Fig. 1.**
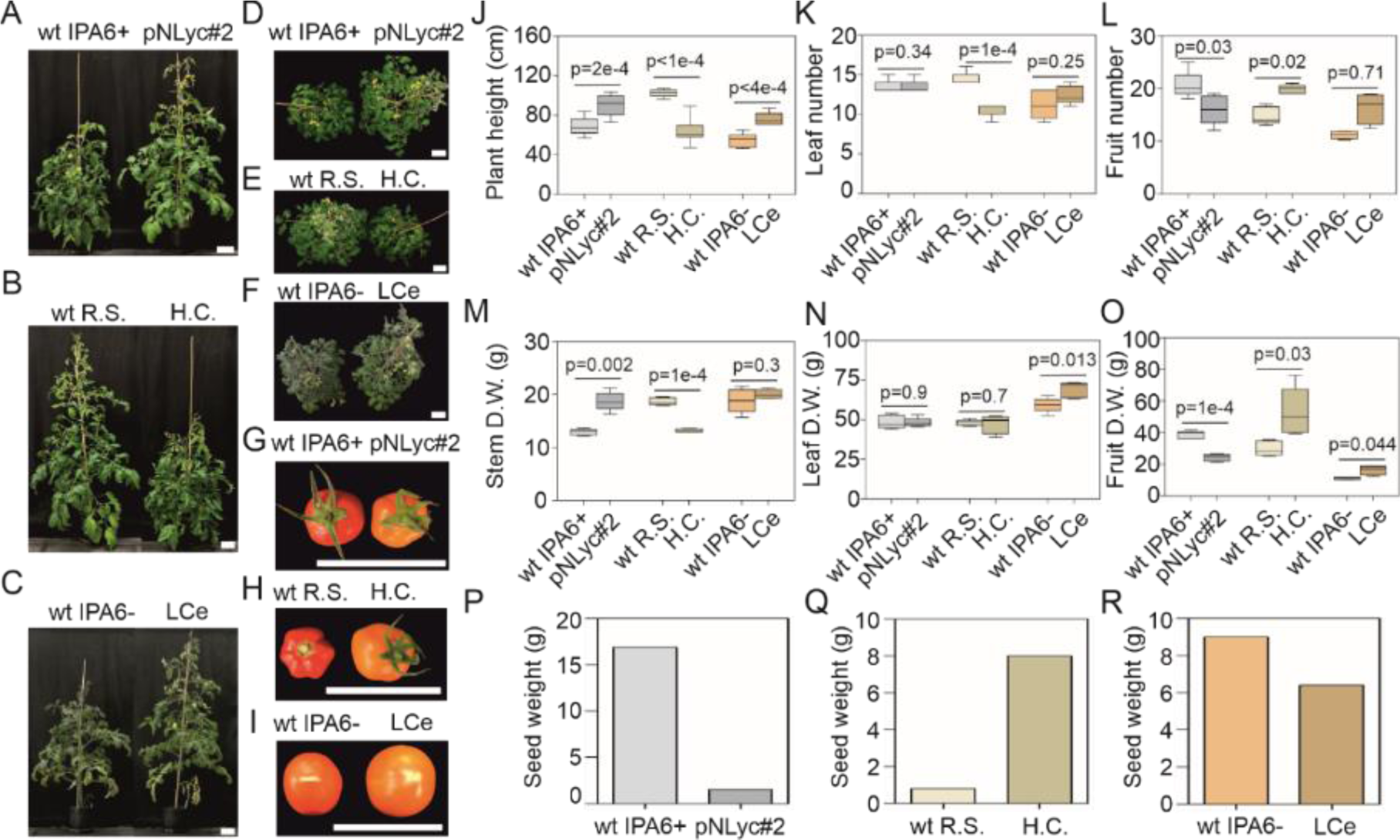
Tomato plant yield under semi-controlled conditions in the greenhouse. **(A-F)** Nine-week-old wild type (IPA6+, R.S., and IPA6-) and transgenic tomato lines (pNLyc#2, H.C., and LCe) grown under greenhouse conditions. **(G-I)** Tomato fruits from 16-week-old wild type and transgenic tomato lines grown under greenhouse conditions (top view). **(J-O)** Plant height, leaf and fruit number, and dry weight biomass (leaf, stem, and fruit) of wild type and transgenic tomato lines. **(P-R)** Seed yield of wild type and transgenic tomato lines grown under greenhouse conditions. Seed production was measured as the total weight of seeds produced by 12 independent tomato plants of each genotype. Five to 10 biological replicates were used (**J-O**). Unpaired two-tailed Student t-test was performed to compare transgenic lines with the wild type. wt: wild type; R.S.: Red Setter; H.C.: high carotene; LCe: lycopene β-cyclase from *Erwinia*. Scale bar: 10 cm.

**Table 1.**
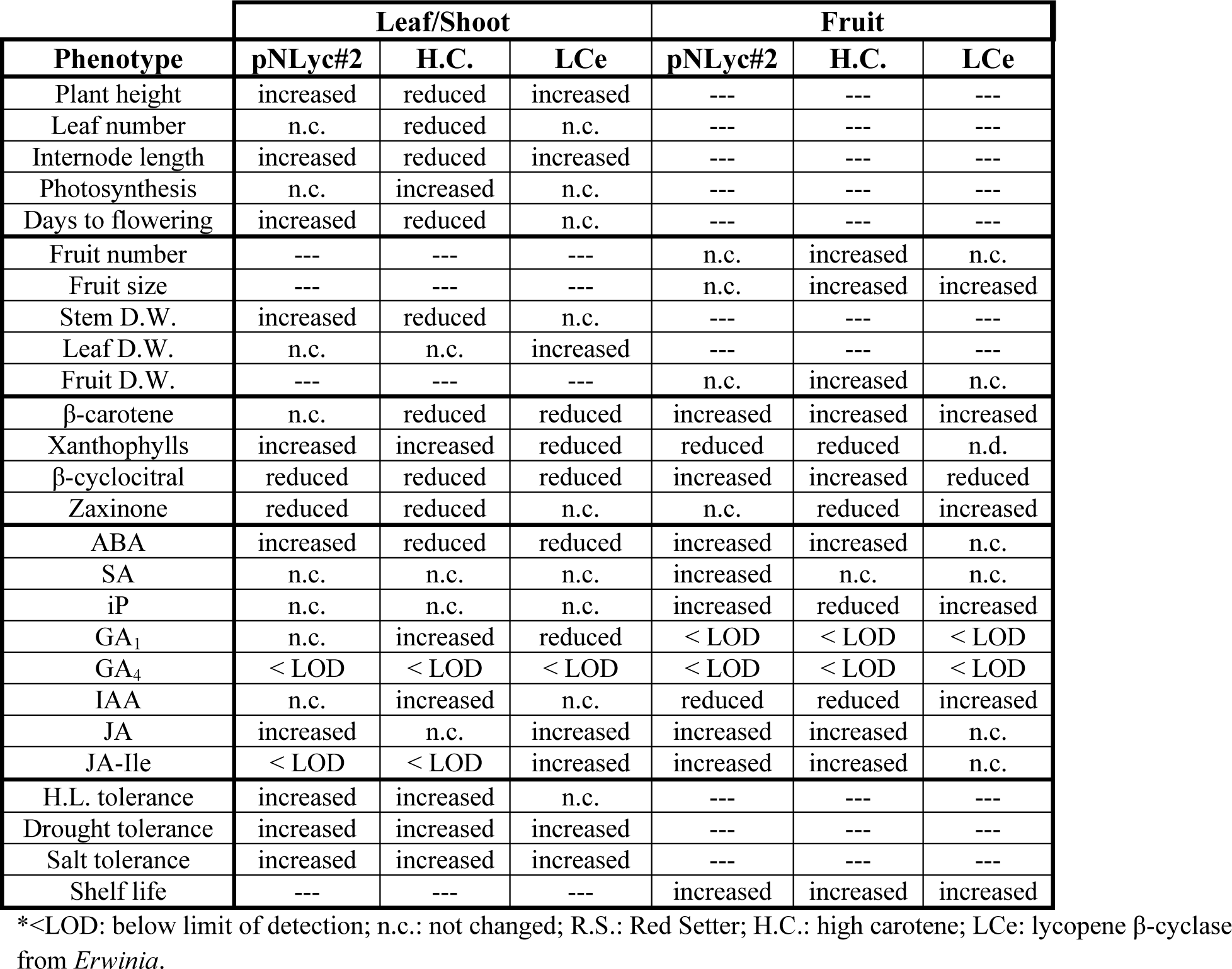
Summary of phenotypic and molecular changes in leaves and fruits of transgenic LCYB-expressing tomato lines

Plant biomass was assessed in all the lines to determine plant productivity. Interestingly, the different *LCYB* transgenic lines showed different biomass partitioning when comparing leaves, stem, and fruit (**Fig. 1M-O**). For instance, the transplastomic pNLyc#2 showed a clear increase in plant height (∼30%) and stem biomass (45%), but no changes in leaf biomass or leaf number (**Fig. 1J, K, M, N**). In addition, fruit biomass (37%) and fruit number were reduced, although the fruit size observed in pNLyc#2 was similar to the wild type (**Fig. 1L**). By contrast, the H.C. line showed reduced plant height (40%) and stem biomass (30%), but no changes in leaf biomass (**Fig. 1J, M, N**). Interestingly, the H.C. line displayed a reduced number of leaves compared to the wild type (**Fig. 1K**). In addition, the H.C. fruit biomass was increased by 77% compared to the wild type R.S. (**Fig. 1O**), in line with the increased fruit number and size displayed by this genotype (**Fig. 1H, L**). The LCe transplastomic line showed increased plant height (∼20%) and leaf biomass (17%), but no significant changes in stem biomass (**Fig. 1J, M, N**). Its fruit biomass was increased up to 45% relative to the wild type IPA6- (**Fig. 1O**). In this line, the leaf and fruit number remained the same as in the wild type (**Fig. 1K, L**). Seed production in pNLyc#2 and LCe transplastomic lines was lower than in their wild types, while H.C. seed production was approximately 1000% higher than in its respective wild type (**Fig. 1P-R**). Biomass quantification in plants grown under fully controlled and uncontrolled conditions showed similar patterns of biomass redistribution (as in the greenhouse) in the different plant tissues (**figs. S3-S4**), but also revealed delayed and accelerated development for the pNLyc#2 and H.C. lines, respectively, while the LCe line showed wild-type-like development (**figs. S4-S5**).

### *LCYB*-overexpressing lines show different carotenoid profiles in leaves and fruit

We sought further insights into the different biomass accumulation patterns in leaves and fruit in the transgenic lines by investigating carotenoid accumulation in both organs, since an altered carotenoid content might affect hormone content and, thereby, plant growth. Transgenic lines expressing plant *LCYB*s showed a reduction in total leaf carotenoid content, with strong decreases in lutein and a lesser decrease in neoxanthin, but strong increases in violaxanthin and zeaxanthin levels. In addition, the H.C. line displayed a slight reduction in β-carotene levels. By contrast, the total carotenoid content in the bacterial *LCYB*-expressing LCe line remained essentially the same as in the wild type, with some slight reductions in β-carotene and zeaxanthin levels in the leaves (**Fig. 2A and fig. S6A, C, E**).

**Fig. 2.**
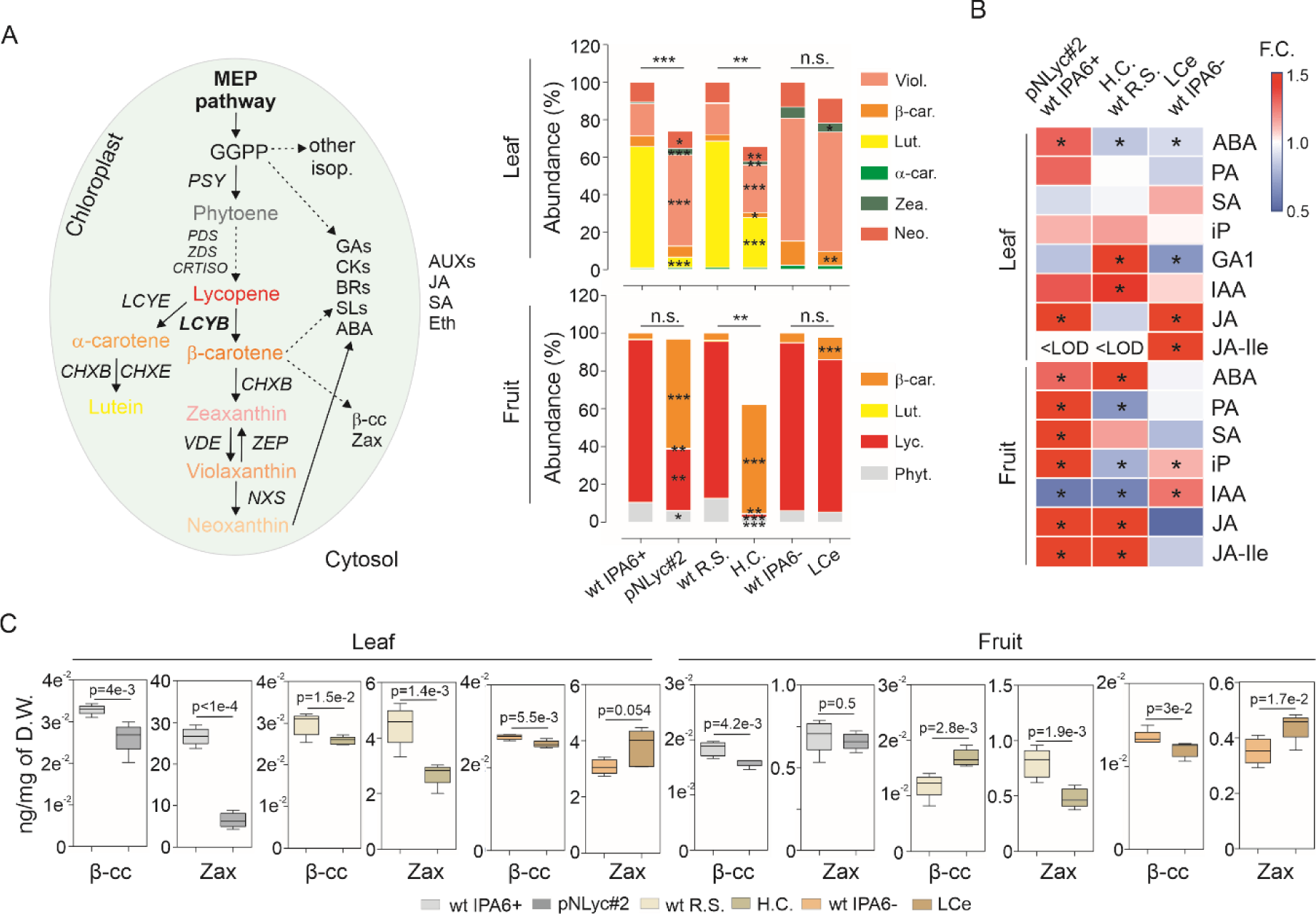
Carotenoid and hormone metabolism in leaf and fruit of *LCYB*-expressing tomato lines. **(A)** Carotenoid pathway (left) and carotenoid composition (right) in leaves and fruits of wild type (IPA6+, R.S., and IPA6-) and *LCYB* transgenic lines (pNLyc#2, H.C., and LCe) grown in the greenhouse. **(B)** Hormone profile in leaves and fruits of wild type and transgenic *LCYB* lines (see **figs. S9-10**). **(C)** Quantification of apocarotenoids with conserved growth-promoting properties (β-cyclocitral/β-cc and zaxinone/Zax) in leaves and fruits (see **figs. S11-15**). Leaf samples were collected from the 5^th^ leaf of each of the five biological replicates used per line (six-week-old plants). Fully ripened fruits were collected from 16-week-old tomato plants (from five different biological replicates, each biological replicate comprising a pool of 3 fruits). Unpaired two-tailed Student t-test was performed to compare transgenic lines with the wild type. In **A**, *: p < 0.05, **: p < 0.005 ***: p < 0.0005; in **B**, *: p < 0.05. wt: wild type; R.S.: Red Setter; H.C.: high carotene; LCe: lycopene β-cyclase from *Erwinia*; LOD: limit of detection; F.C.: fold change. Viol: violaxanthin; car: carotene; Zea: zeaxanthin; Neo: neoxanthin; Lyc: lycopene; Phyt: phytoene; Lut: lutein. ABA: abscisic acid; PA: phaseic acid; IAA: indole acetic acid; iP: isopentenyladenine; GA: gibberellins; SA: salicylic acid; JA: Jasmonic acid; Ile: isoleucine; Eth.: ethylene; isop.: isoprenoids. PSY: phytoene synthase; PDS: phytoene desaturase; ZDS: z-carotene desaturase; CRTISO: carotene isomerase; CHX: carotenoid hydroxylase; VDE: violaxanthin de-epoxidase; ZEP: zeaxanthin epoxidase; NXS: neoxanthin synthase.

In the fruit, the total carotenoid content in the transplastomic pNLyc#2 and LCe lines was unchanged, while the total carotenoid content was reduced in the H.C. line. Transgenic lines expressing plant *LCYB*s showed a strong accumulation of fruit β-carotene and strong reductions in lycopene, lutein, and phytoene, while an increase in β-carotene was only observed for the bacterial *LCYB*-expressing LCe line (**Fig. 2A and fig. S6B, D, F**). In addition, carotenoid-rich crystal structures were observed by confocal microscopy in the fruit of the transgenic lines (**fig. S7**). Due to the possibility that other isoprenoid pathways might have been affected by *LCYB* expression (Moreno *et al*., 2020), we also determined the chlorophyll and tocopherol (vitamin E) content in the leaves and fruit (**fig. S8**). Chlorophyll contents remain unchanged in the pNLyc#2 and H.C. lines (with the exception of a slight reduction in chlorophyll b in the H.C. line), while γ- and α-tocopherol contents were increased. The LCe line showed a reduction in α-tocopherol (**fig. S8**). By contrast, the tocopherol content (α, δ, and γ-tocopherol) increased strongly in fruit of the pNLyc#2 line, while remaining unaltered in the H.C. and LCe lines (**fig. S8**).

### Hormone metabolism is altered in *LCYB*-overexpressing lines

Altered β-carotene accumulation might influence the content of β-carotene-derived and/or isoprenoid-derived hormones (e.g., ABA and Gas, respectively), thereby influencing plant growth and development. Therefore, we profiled the plant hormones to gain further insights into their contribution to the observed growth phenotype. The lines were characterized by significant increases in ABA and jasmonic acid (JA) for pNLyc#2; ABA reduction and GA_1_ and IAA increments for H.C.; and ABA and GA_1_ reductions and JA and JA-Ile increments in LCe in leaves (**Fig. 2B**). By contrast, stronger significant changes in hormone content were found in fruit. ABA, JA, and JA-Ile were increased, while indole acetic acid (IAA), the most bioactive auxin (Aux), was reduced in both the pNLyc#2 and H.C. lines but increased in the LCe line (**Fig. 2B**). In addition, SA was increased only in the pNLyc#2 line, whereas isopentenyladenine (iP), an active cytokinin (CK), was increased in the pNLyc#2 and LCe lines (**Fig. 2B**). Phaseic acid, a bioactive ABA catabolite, showed increased and reduced contents in the pNLyc#2 and H.C. lines, respectively. Intermediates of the ABA, GA, Aux, CKs, and JA metabolic pathways were also differentially affected in leaves and fruit (**fig. S9-10**).

### Effects of carotenoid accumulation on apocarotenoid metabolism in leaves and fruit

β-carotene and xanthophylls are the main precursors of non-hydroxylated and hydroxylated apocarotenoids, respectively. Growth-promoting and signaling properties of some apocarotenoids (e.g., β-cyclocitral and zaxinone) have been reported in rice, tomato, and Arabidopsis (Dickinson et al., 2019; Wang et al., 2019). These previous findings and the altered pigment content observed in the leaves and fruit of the transgenic lines led us to profile apocarotenoid species in order to determine their contribution to the observed phenotypes (**Fig. 2C and fig. S11-15**). In leaves, non-hydroxylated apocarotenoids showed few increases or wild-type-like accumulation (**fig. S12**), in line with the wild-type-like β-carotene content in the transgenic lines. By contrast, hydroxylated apocarotenoids showed strong reductions due to a strong decrease in lutein content (**fig. S13**). The non-hydroxylated apocarotenoids in fruit showed a strong and significant accumulation (due to enhanced β-carotene content; **fig. S14**), while the hydroxylated apocarotenoids exhibited strong reductions due to the lower lutein content in the fruit (**fig. S14**). Growth regulators, such as β-cc and Zax, were mainly found at reduced levels in the leaves and fruit (**Fig. 2C**). Other apocarotenoids with biological activity, such as β-ionone, showed enhanced accumulation in the fruit (**fig. S11**).

### Primary metabolites and lipid metabolism are altered in leaves and fruit of *LCYB*-expressing lines

The strong changes in pigment, hormone, and apocarotenoid contents led us to investigate the impact of these changes on other metabolic pathways. GC-MS metabolite profiling showed significant changes in sucrose and its derivatives (e.g., fructose, galactinol, raffinose), glycolytic intermediates (e.g., glucose, G6P, Fru6P) and TCA cycle intermediates (e.g., malate and fumarate) in the leaves and fruit of the transgenic lines (**Fig. 3B; fig. S16**). These changes were reflected, for instance, in changes in G6P-derived compounds (e.g., trehalose, maltotriose, maltose, *myo*-inositol, and erythritol) and amino acids derived from glycerate (e.g., O-acetylserine [OAS]), pyruvate (e.g., valine, alanine, leucine), shikimate (e.g., phenylalanine and tryptophan), malate (e.g., aspartic acid, asparagine, β-alanine, and methionine), and 2-oxoglutarate (e.g., glutamic acid, glutamine, GABA, and ornithine) (**Fig. 3B**). In addition, due to the structural function of carotenoids (β-carotene and xanthophylls) in membrane composition, together with lipids, we determined the lipid composition in leaves and fruit. Lipid profiling revealed no significant differences in the leaves, while marked significant differences were observed, mainly for structural lipids, in the fruit of pNLyc#2 and H.C. lines (**Fig. 3B; fig. S17**). In the fruit of pNLyc#2, a total of 17 galactolipids (GLs) (e.g., mono- and di-galactosyldiacylglycerol, [MGDG and DGDG, respectively]) and 32 phospholipids (PLs) (e.g., phosphatidylcholine [PC], phosphatidylethanolamine [PE], phosphatidylglycerol [PG], and phosphatidylserine [PS]) exhibited significant changes in their ratio abundances (**Fig. 3B)**, with levels of nine GLs reduced and eight increased, while the trend for PLs differed, where abundance rations were reduced for seven PLs and increased for 25 PLs. The general trend for sulfolipids (SLs) (e.g., sulfoquinovosyl diacylglycerol [SQDG]) and di- and tri-acylglycerols (DAGs/TAGs) was a reduced abundance, with the exception of two SL species (**Fig. 3B**). By contrast, in the H.C. line, most of the lipid species that showed significantly different levels displayed a reduced ratio abundance, with a few exceptions (e.g., two DAGs, four TAGs, one PC, and two PEs) that showed increased content (**Fig. 3B**).

**Fig. 3.**
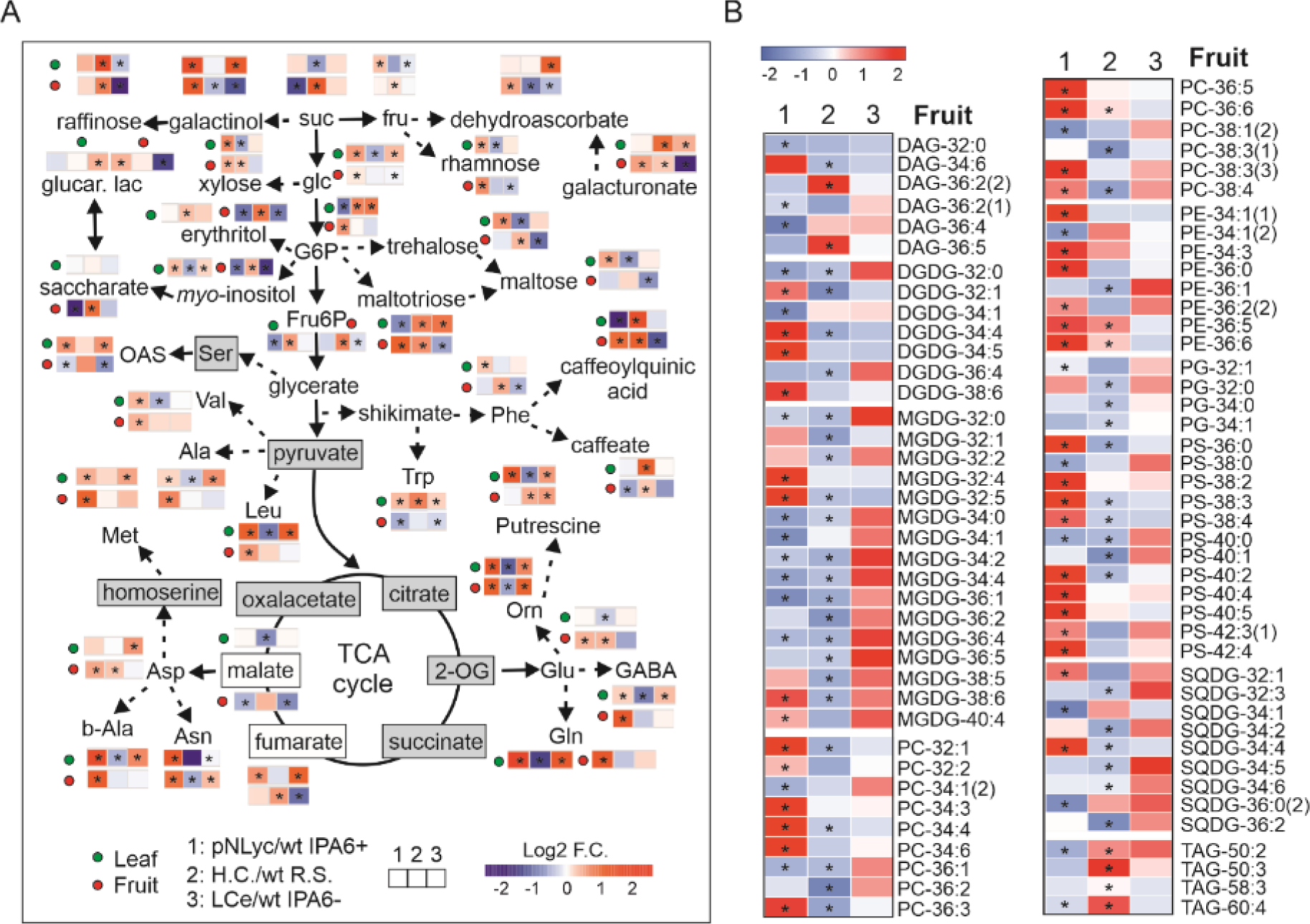
Metabolic reshaping in leaves and fruits by *LCYB* expression in tomato. **(A)** Primary metabolite profiling in leaves and fruits of wild type (IPA6+, R.S., and IPA6-) and *LCYB* transgenic tomato lines (pNLyc#2, H.C., and LCe). A non-paired two-tailed Student t-test was performed to compare each transgenic line with their wild type (p<0.05; *n*=5 biological replicates). **(B)** Lipid profile in fruits of *LCYB* transgenic tomato lines. The lipid profile in leaves is reported; however, no significant changes were observed (**fig. S17**). Wilcoxon’s test was performed to compare transgenic lines with their wild types (p<0.05; *n*=5 biological replicates). Changes are shown as the log2 fold change between the transgenic lines and their respective wild type controls (for more details see **fig. S16-17**). Asterisks represent significant changes. OG: oxoglutarate; orn: ornithine; GABA: gamma aminobutyric acid; suc: sucrose; fru: fructose; glc: glucose; G6P: glucose-6-phosphate; Fru6P: fructose-6-phosphate; OAS: o-acetylserine; glucar. lac: glucarate-1,4-lactone; DAG: diacylglycerol; DGDG: di-galactosyldiacylglycerol; MGDG: mono-galactosyldiacylglycerol; PC: phosphatidylcholine; PE: phosphatidylethanolamine; PG: phosphatidylglycerol; PS: phosphatidylserine; SQDG: sulfoquinovosyl diacylglycerol; TAG: triacylglycerol.

### Photosynthetic parameters are influenced by carotenoid accumulation and plant architectural changes in tomato *LCYB*-expressing lines

The changes in plant growth and architecture induced by modifications in pigment and hormone contents prompted subsequent analysis of several photosynthetic parameters. Photosynthetic measurements were performed in tomato plants (49 days old) grown under greenhouse conditions (**fig. S18**). CO_2_ assimilation was significantly increased for the H.C. line, relative to its wild type, whereas the transplastomic lines were the same as their wild types (**Fig. 4A**). Despite some unaltered photosynthetic parameters, the ΦPSII, which reflects plant fitness, was increased in all the lines (**Fig. 4B**). Interestingly, NPQ(T) was reduced in the H.C. line but was unaltered in the transplastomic lines, in agreement with the observed ΦNPQ (**Fig. 4C and fig. S18H**). Conductance was also reduced in the pNLyc#2 line and increased in the H.C. and LCe lines (**fig. S18F**). The rETR was unchanged in the transplastomic pNLyc#2 and LCe lines but was increase in the H.C. line (**fig. S18G**). These results suggest that the nuclear H.C. line is the one with the most enhanced photosynthetic efficiency, despite its smaller shoot size.

**Fig. 4.**
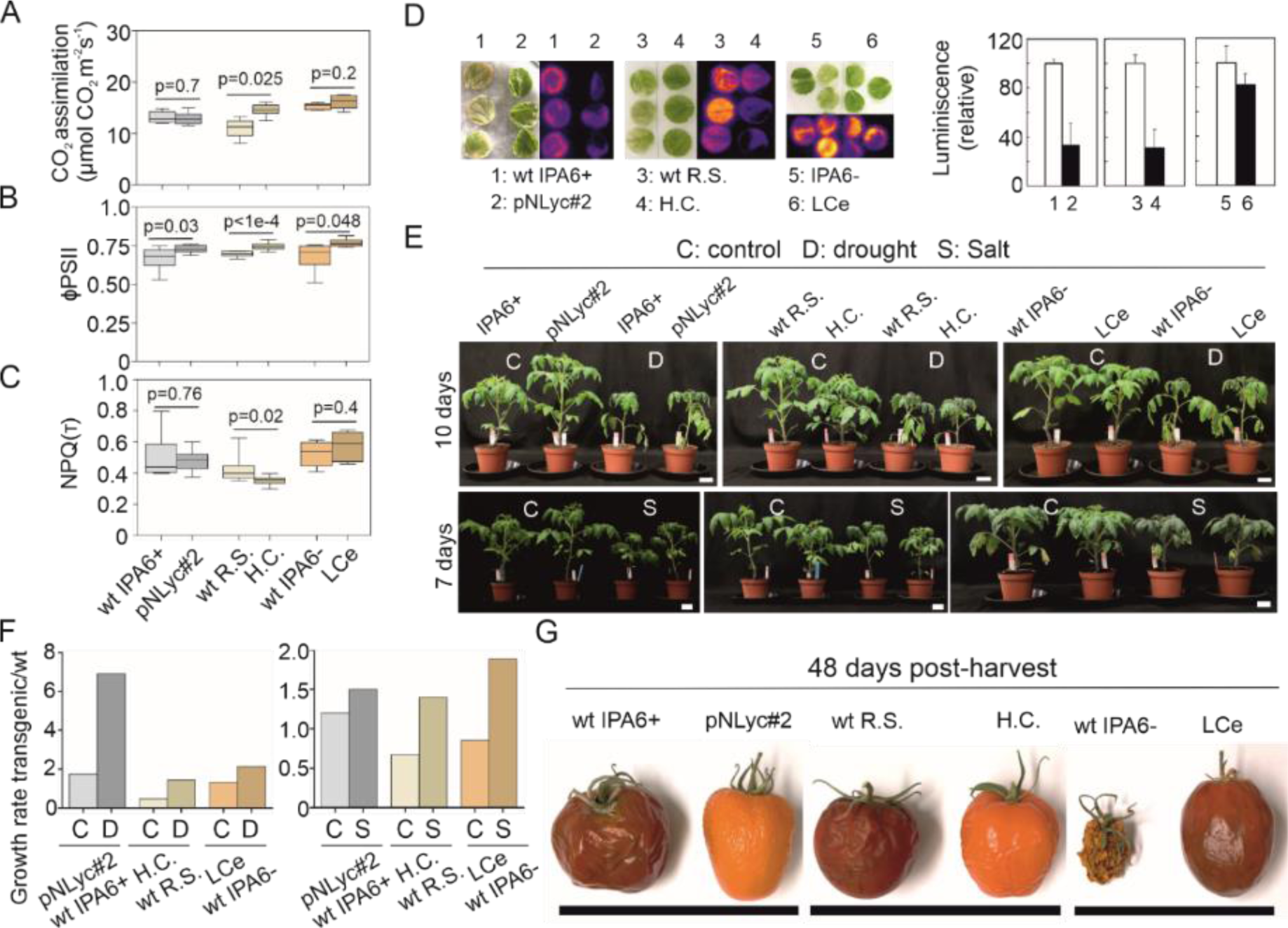
Photosynthetic parameters, stress tolerance, and shelf life of transgenic *LCYB* tomato lines. **(A)** CO_2_ assimilation. **(B)** ΦPSII. **(C)** Total non-photochemical quenching (NPQT). CO_2_ assimilation was measured with a Li-Cor instrument and ΦPSII and NPQT with a MultiSpec instrument (Photosync). Photosynthetic parameters were measured from leaves of seven-week-old wild type (IPA6+, R.S., and IPA6-) and transgenic (pNLyc#2, H.C., and LCe) tomato lines grown under greenhouse conditions. All measurements, and especially NPQT, were performed without a dark adaptation period, as described in Tietz et al. *(31)*. Five to 12 biological replicates were used for each photosynthetic measurement. **(D)** Lipid peroxidation imaging and quantification of tomato leaf discs (six-week-old plants) exposed to a light intensity of 2000 μmol photons m^-2^ s^-1^ and a temperature of 7°C degrees. **(E)** Water deficit and salt treatments in three-week-old wild type and transgenic lines (*n*=5-6) grown in the greenhouse in 13 cm pots (see material and methods). Plant height was recorded before and after water deficit and salt treatments. **(F)** Growth rate (plant height) ratio between transgenic lines and their respective wild type controls. Plant height was measured before (0 days) and after stress onset (10 days for water deficit and seven days for salt treatments) and the growth rate was calculated under control and stress conditions. **(G)** Tomato shelf life in wild type and transgenic tomato fruits. Tomato fruits from wild type and transgenic lines were harvested from 15-week-old tomato plants. Shelf life was recorded at 48 days post-harvest (see **fig. S20** for other time points). A non-paired two-tailed Student t-test was performed to compare transgenic lines with the wild type. wt: wild type; R.S.: Red Setter; H.C.: high carotene; LCe: lycopene β-cyclase from *Erwinia*.

### *LCYB*-expressing lines show enhanced abiotic stress tolerance and shelf life

The increases in xanthophyll and hormone contents were further assessed, given their functions in photoprotection and stress tolerance, by exposing the transgenic lines to abiotic stress. Leaves of the pNLyc#2 and H.C. transgenic lines, which had higher xanthophyll content, showed high light tolerance, as measured by the luminescence produced by the accumulation of lipid peroxides (**Fig. 4D**). The LCe line showed no significant increase in high light tolerance (**Fig. 4D**). In addition, all the transgenic lines showed higher growth rates when exposed to either water deficit or salinity treatments (for 10 and seven days, respectively) when compared to their wild type counterparts (**Fig. 4E-F and fig. S19**). An extended fruit shelf-life has previously been reported in tomato and other fruit due to enhanced ABA content or to the content of other primary metabolites (e.g., putrescine), so we also examined fruit shelf-life in the transgenic lines. All transgenic lines showed enhanced shelf-life at different time points after harvest when compared to their respective wild types (**Fig. 4G and fig. S20**).

## DISCUSSION

The tomato is one of the most important fruit and vegetable crops worldwide, but its productivity is affected by several abiotic stresses that have deleterious effects on fruit number and size, as well as on fruit quality (Gerszberg and Hnatuszko-Konka, 2017). In the present study, we have demonstrated that *LCYB* expression has beneficial effects on tomato plant fitness, stress tolerance, and biomass, regardless of the *LCYB* genetic origin, tomato cultivar, or genetic transformation strategy (**Table S1**). However, the mechanisms by which the introduction of a *LCYB* gene modulates plant growth and development, photosynthetic efficiency, and stress tolerance remain unresolved. LCYB catalyzes the conversion of lycopene to β-carotene, a step previously characterized as a metabolic hot spot in tobacco (Kossler et al., 2021; Moreno *et al*., 2020). The metabolic hot spot focused on β-carotene reflects its multiple functions in several molecular and physiological processes (e.g., photosynthesis, oxidative stress). In addition, β-carotene serve as precursor of xanthophylls (photoprotection), hormones (growth, development, and stress response), and growth regulators (**Fig. 5A**). Thus, changes in carotenoid content could directly influence photosynthesis, antioxidant properties, and pigment content, while also indirectly influencing hormone and apocarotenoid content (ABA, SLs, β-cc) and, consequently, plant growth, development, and stress responses (Al-Babili and Bouwmeester, 2015; Nambara and Marion-Poll, 2005; Wang *et al*., 2019).

**Fig. 5.**
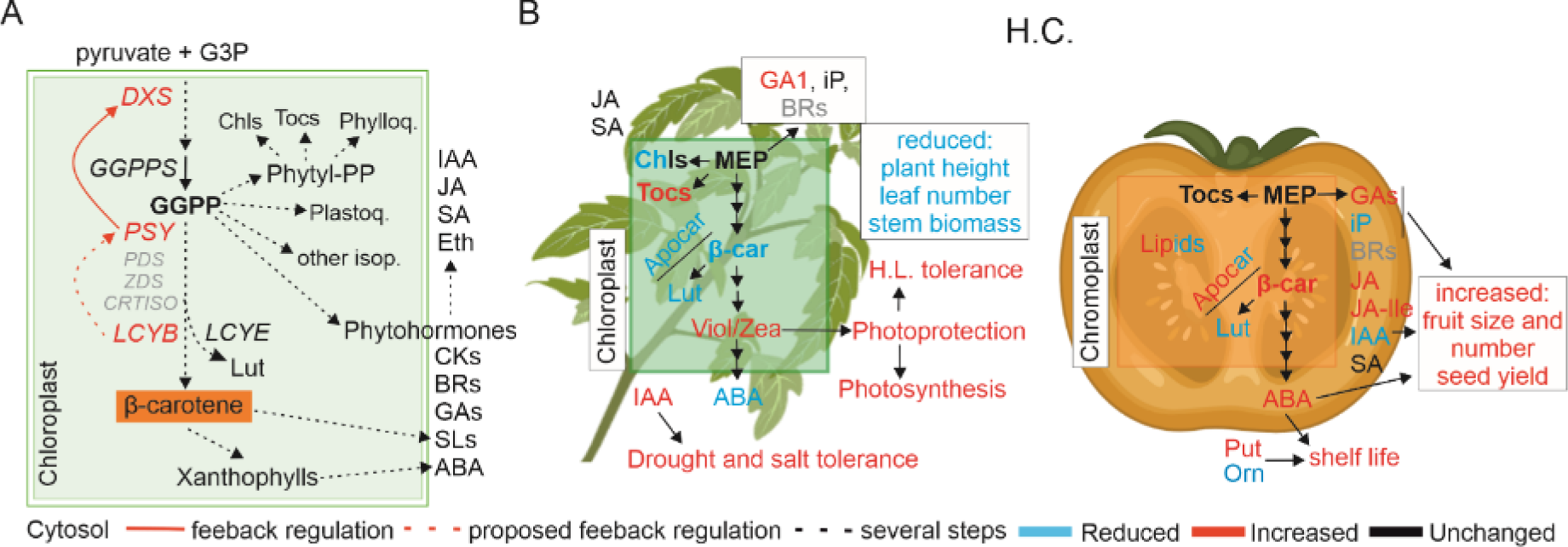
Proposed model for *LCYB*-mediated plant fitness enhancement. **(A)** Schematic representation of isoprenoid pathways connected by the common precursor GGPP. Conversion of lycopene into β-carotene represents a major key regulatory point in the branching of the carotenoid pathway. The greater β-carotene production is used for greater production of xanthophylls (photoprotection) and hormone synthesis (modulation of plant growth, development, and stress tolerance). Feedback regulation between *LCYB*, *PSY*, and *DXS* might be controlling the production of GGPP and therefore influencing the content of other isoprenoids (e.g., GAs, tocopherols, and chlorophylls). **(B)** Metabolic and physiological changes in leaves (left side) and fruits (right side) of the high carotene (H.C.) tomato transgenic line showing the influence on yield, stress tolerance, photosynthetic efficiency, pro-vitamin A content, and fruit shelf life (for comparison with transplastomic lines see **fig. S21**). Increases (red), reductions (blue), no changes (black), or compounds under the detection limit by the hormonomics approach (grey), are shown. Metabolites (e.g., carotenoids, apocarotenoids, hormones, lipids) with different accumulation profiles (increases and decreases in different metabolites) are shown both in red and blue. Put: putrescine; Orn: ornithine; Lut: lutein; β-car: β-carotene; Tocs: tocopherols; Chls: chlorophylls; Apocar: apocarotenoids; GAs: gibberellins; Viol: violaxanthin; Zea: zeaxanthin; BRs: brassinosteroids; iP: isopentenyladenine.

Feedback mechanisms between carotenoids, methylerythritol phosphate (MEP), and ABA pathways, can also influence carotenoid accumulation in maize, rice, Arabidopsis, and tomato (Bai et al., 2009; Beyer et al., 2002; Qin et al., 2007; Romer et al., 2000). Enhanced *PSY* expression in etiolated Arabidopsis seedlings also resulted in enhanced carotenoid levels via post-translational accumulation of *DXS* mRNA, which stimulated the supply of MEP substrates (Rodriguez-Villalon et al., 2009b; a). Thus, any alteration in the expression of a carotenogenic gene can impact the expression of other carotenoid genes, as well as key genes from other isoprenoid pathways (e.g., *DXS*, *GA20ox, CHL*), as observed in *DcLCYB1* tobacco lines (Moreno *et al*., 2020). This reflects a close interconnection between the isoprenoid pathways (**Fig. 5A**) and suggests that any disturbance in the metabolic flux of a particular isoprenoid pathway (e.g., carotenoid pathway) may affect other plastidial isoprenoid-related pathways. Notably, isoprenoids are also the precursors of gibberellins (GAs), brassinosteroids, and cytokinins (CKs), so any disturbance in the isoprenoid flux might influence hormone contents, with subsequent impacts on plant growth, development, and stress tolerance (Gudesblat and Russinova, 2011; Ha et al., 2012; Hedden and Phillips, 2000; Krishna, 2003; Schaller et al., 2015; Tran et al., 2007). In fact, the transgenic tomato lines analyzed here are evidence of carotenoids as a metabolic hot spot (**Fig. 5**) because, despite the differences in their genetic background, these tomato lines universally displayed changes in carotenoids, apocarotenoids, and hormone contents (**Fig. 2**) that resulted in altered growth regulation and biomass partitioning in different tissues (**Fig. 1 and figs. S1-4**). These changes were furthermore reflected in plant biomass accumulation, resilience to abiotic stresses, and crop productivity (**Fig. 1 and Fig. 4D-F**).

The hormonal changes and their effects on primary metabolism can explain the changes in biomass accumulation and stress tolerance (Moreno *et al*., 2021; Moreno *et al*., 2020; Sheyhakinia et al., 2020; Yoshida et al., 2014). For instance, gibberellins (GAs) control many aspects of growth (e.g., plant height, internode length) and plant development. Bioactive GAs (GA_4_ and GA_1_) function as key players in plant growth and development in Arabidopsis, tobacco, and rice, with GA_4_ showing the highest bioactivity (Cowling et al., 1998; Gallego-Giraldo et al., 2008; Talon et al., 1990; Ueguchi-Tanaka et al., 2007). Both the bioactive GAs are produced from GA_12_ by the non-13-hydroxylation (GA_4_) and the early-13-hydroxylation (GA_1_) pathways (Magome et al., 2013). Interestingly, manipulation of GA biosynthetic genes (e.g., *COPALYL SYNTHASE*, *GA3oxidase 1*, *GA20oxidase 1*) in Arabidopsis, tobacco, and rice, showed opposite GA_4_ and GA_1_ accumulation patterns (Fleet et al., 2003; Gallego-Giraldo *et al*., 2008; Magome *et al*., 2013). In our lines, the longer stems and internode lengths (**Fig. 1A, C, J, and fig. S18A, D**) in the transplastomic lines suggest an enhanced GA_4_ content. However, the GA_4_ level was below the detection limit in the material we profiled in our study, although we detected a reduction in GA_1_, which could potentially reflect an increase in GA_4_ in the transplastomic lines. By contrast, the shorter stem and internodes, together with enhanced GA_1_ content, in the H.C. line suggest a possibly decreased GA_4_ content (**Fig. 1B, J, and fig. S18A, D**).

The reduced-growth phenotype is in line with the reduced plant size previously reported in ABA-deficient mutants of tomato (Nitsch et al., 2012). However, a similar ABA reduction in LCe, which shows an opposite phenotype to H.C. (longer stem and internodes), suggests that the interaction between GA_4_ and ABA might direct plant height, as previously observed in *DcLCYB1* tobacco lines (Moreno *et al*., 2020). In addition, reductions in β-cyclocitral and/or zaxinone in the transplastomic lines (**Fig. 2C**) suggest that they are not involved in the observed growth phenotype, while reductions in both metabolites might contribute to the smaller growth phenotype observed in the H.C. line (**Fig. 1**). The enhanced ABA and JA (p=1.1e^-3^ and p=4e^-3^; **Fig. 2B and fig. S9**) contents in pNLyc#2 are likely responsible for its salt and drought tolerance (**Fig. 4E, F, and fig. S19**), as previously shown in Arabidopsis and tobacco (Kazan, 2015; Moreno *et al*., 2021; Moreno *et al*., 2020; Yoshida *et al*., 2014).

An enhanced ABA content may have caused stomatal closure, as reflected in the observed reduction in stomatal conductance (**fig. S18F**). This reduction would conceivably impede an enhancement of photosynthetic efficiency (**Fig. 4A-C, and fig. S18F-I**). By contrast, the H.C. and LCe lines displayed a slightly reduced ABA content and enhanced conductance; however, only the H.C. line showed enhanced photosynthetic efficiency (higher CO_2_ assimilation, rETR, and ΦPSII; **Fig. 4A-C, and fig. S18F-I**). Although these lines showed reduced ABA content, they both showed enhanced salt and drought tolerance, suggesting the participation of an ABA-independent pathway. In fact, JA/JA-Ile are involved in salt and drought tolerance in Arabidopsis and rice (Hazman et al., 2019; Kazan, 2015). Increases in JA and JA-Ile (p=0.05 and p<1e^-4^; **Fig. 2B and fig. S9**) in the LCe line supported the higher drought and salt tolerance observed in this line. However, the H.C. line showed reductions in ABA and no changes in JA, but a significant increase in IAA (p=0.03; **Fig. 2B and fig. S9**).

IAA has been reported to enhance salt and drought tolerance in white clover, Arabidopsis, and rice (Shani et al., 2017; Sharma et al., 2013; Shi et al., 2014; Zhang et al., 2020), supporting its enhanced tolerance to these abiotic stresses (**Fig. 4E, F, and fig. S19**). In addition, several osmoprotectants, which are neutral molecules that help the organisms to persist during severe osmotic stress (Singh et al., 2015), were enhanced in the transgenic lines (**Fig. 3A**). Increased ABA and JA contents were previously reported to enhance the synthesis of osmoprotectants (e.g., sugars, polyamines) under abiotic stress conditions to counteract harmful effects (Alcazar et al., 2006; Sheyhakinia *et al*., 2020; Toumi et al., 2010; Wang et al., 2020). In line with this evidence, increases in sugars (raffinose, fructose, G6P, glucose, trehalose), sugar alcohols (*myo*-inositol, erythritol) and polyamines (putrescine) in leaves can also contribute to enhanced stress tolerance in our transgenic lines (**Fig. 3A, Fig. 4E, F, and fig. S19; Table S1**).

The increased xanthophyll content in leaves could further enhance photoprotection and therefore impart high light tolerance (pNLyc#2 and H.C.; **Fig. 4D**). In the fruit, stronger increases in β-carotene content caused stronger changes in hormone content, thereby impacting fruit dry matter (up to 67–77% in semi-controlled and uncontrolled conditions, respectively), size, and number, as well as seed production (**Fig. 1, Fig. 2A, B, and fig. S4; Table S1**), making the fruit rich in pro-vitamin A and enhancing its nutritional value. Fruit growth is influenced by CKs, Aux, GA, and ABA (Quinet et al., 2019). Transgenic fruit differentially accumulate IAA, iP, ABA, and GA intermediates, suggesting that their interaction may have led to the observed fruit growth phenotypes (**Fig. 1G-I, and fig. S2B, C, E, F, H, I, fig. S4C, D, G, H, K, L, fig. S5, fig. S8B, fig. S10**).

Unfortunately, GA_1_, which was reported to be the most bioactive GA influencing fruit growth (Garcia-Hurtado et al., 2012), was under the detection limit in fruit in our experiments, but its content might explain the large increase in fruit size in the H.C. line. Furthermore, changes in the hormonal network might confer additional advantages to the shoots or fruit. Recently, Diretto et al. showed that the enhanced shelf-life of *LCYB*-expressing tomato lines was due to increased ABA content and its negative impact on ethylene content (Diretto et al., 2020). Increased ABA content in the pNLyc#2 and H.C. lines conferred longer fruit shelf-life compared to the wild type (**Fig. 4G and fig. S20**). However, in the LCe line, which also showed enhanced shelf-life, the ABA content was unchanged, suggesting that shelf-life might be controlled by other factors. Indeed, polyamines (e.g., spermidine, putrescine) are known anti-senescence agents which increase fruit firmness, delay ethylene emission and the climacteric respiratory burst, and induce mechanical stress resistance (Valero et al., 2002). The highest ornithine and putrescine content (p<0.05) was observed in the LCe line, and this could contribute to the enhanced shelf-life observed in the fruit of this line (**Fig. 3A; Table S1**).

Accumulation of sugars and derivatives (e.g., raffinose, galactinol, *myo*-inositol, and trehalose) and amino acids (e.g., Val, Asp, Asn, Thr, Glu, Gln, and Ala) in fruit were reported to confer tolerance to chilling injury and resistance to pathogens and several postharvest stress conditions (Bang et al., 2019; Farcuh et al., 2018; Lauxmann et al., 2014; Luengwilai et al., 2018). Accumulation of these metabolites would be expected to confer valuable post-harvest traits to our tomatoes apart from the enhanced shelf-life and their higher pro-vitamin A content.

The use of transgenic tomato lines with different cultivar and genetic backgrounds allowed us to demonstrate that i) *LCYB* overexpression can be used to modulate growth (different biomass partitioning between leaf and fruit) and fruit yield in a crop, and ii) the positive growth regulatory effect conferred by the carrot *DcLCYB1* gene in tobacco (Moreno *et al*., 2020) can be also conferred by other LCYBs (e.g., tomato, daffodil, and bacteria) in leaves and/or fruit. However, the different genetic origins of the chosen *LCYB* genes also introduced specific changes in each line (**Table 1; fig. S21**). Therefore, the selection of the transgene should be carefully analyzed before using it for biotechnological purposes.

In conclusion, while some of the differences at the phenotypic (e.g., biomass partitioning; **Table 1; fig. S21**) and molecular levels observed in the transgenic lines might reside in the different cultivars, transformation methods, and *LCYB* genetic origins (**Table S1**), many similarities can be explained by the modulation of molecular processes, such as carotenoid and hormone accumulations (see above; **fig. S21**). Despite the observed specific changes in carotenoid, hormone, and metabolite accumulation in leaves and fruit of the transgenic lines (**Table 1; Table S2-3 and fig. S21**), the similar responses in these lines can be attributed to changes in specific hormones (salt and drought tolerance are most likely conferred by increases in ABA and JA for pNLyc#2, IAA for H.C., and JA and JA-Ile for LCe; **Table 1; fig. S21**) and/or metabolites (e.g., putrescine-enhanced shelf-life). However, other observed contrasting phenotypes (e.g., plant height and seed yield) were probably caused by specific interactions between hormones and/or their ratios, as well as the connection between carotenoids and other non-isoprenoid hormones (e.g., IAA), and these remain to be investigated. Nevertheless, modulation of the content of main components of the hormonal network in each transgenic line resulted in enhanced abiotic stress tolerance, extended fruit shelf life, and increased biomass (favoring shoot and/or fruit in the different lines), along with the enhanced nutritional value conferred by the higher β-carotene content in the fruit (**Table 1; fig. S21**). All these features are highly desirable traits for crop improvement (especially stress tolerance and higher biomass/yield) considering the worldwide climate change and its consequences for food crop production. This type of bioengineering is a promising strategy that can be exported to cereal crops (e.g., rice) that, in general, do not accumulate high levels of carotenoids but whose yield must be greatly increased by 2050.

## METHODS

### Plant material and growth conditions

Tomato wild type (*S. lycopersicum* cvs. IPA6+/lutein, IPA6-/without lutein and isogenic Red Setter/R.S.), transplastomic (pNLyc#2 and LCe), and nuclear (high carotenoid/H.C.) lines (Apel and Bock, 2009; D’Ambrosio et al., 2004; Wurbs et al., 2007) were raised from seeds germinated on soil. The transgenic lines harbor *LCYB* genes from daffodil, tomato, and bacteria (*Erwinia uredovora*). Two of the selected lines were obtained by plastid DNA transformation (pNLyc#2 and LCe) and the other line by Agrobacterium-mediated nuclear DNA transformation (H.C.; **Table S1**). Transplastomic lines expressing the *LCYB* gene from daffodil or *Erwinia uredovora* (pNLyc#2 and LCe, respectively) were generated via plastid transformation using particle bombardment. The homoplasmic state (i.e., the absence of residual copies of the wild-type genome) of ∼22 plants was assessed by subjecting the transgenic plants to double-resistance tests (spectinomycin and streptomycin, 500 mg l^-1^) on synthetic media and by RFLP analysis (Apel and Bock, 2009; Wurbs *et al*., 2007). Due to the homoplasmic state (meaning that plastid DNA was equally modified in all chloroplasts of the transgenic lines) and to the similar phenotype observed in these lines, we selected one line per genotype (pNLyc#2 and LCe) to carry out the experiments described in this work. The H.C. nuclear line (plus other six LCYB transgenic lines) was obtained via *Agrobacterium* transformation. All seven transgenic lines expressing the tomato *LCYB* were confirmed by Southern blot experiments and by the intense orange color in their fruit in comparison to the isogenic Red Setter control. In addition, northern blot and qPCR experiments confirmed higher transcript accumulations in the transgenic lines in leaves and fruit than in the isogenic wild type Red Setter control (D’Ambrosio *et al*., 2004; Giorio et al., 2007). Based on this evidence and the similar phenotype obtained in all nuclear lines, we selected the H.C. line with the highest β-carotene levels for the experiments in this work.

Wild type and transgenic lines were grown side by side, and randomly allocated, in the greenhouse (semi-controlled conditions) under standard conditions (16 h/8 h day/night regime, 450–800 μmol photons m^-2^ s^-1^ combination of artificial light and sunlight, 24 °C, and 65 % relative humidity). Plant height, leaf and fruit number, internode length, and seed yield were recorded. Fully expanded mature source leaves (the 5^th^ leaf) were harvested from six-week-old wild type and transgenic *LCYB* tomato plants (*n*=5) grown in the greenhouse. Fruits were analyzed as five biological replicates from 16-week-old tomato plants. Each biological replicate consisted of a pool of three different fruits from one individual plant.

### Physiological measurements and biomass quantification

The T5 generation wild type (R.S.) and nuclear transformed (H.C.) and wild type (IPA6+ and IPA6) and T3 transplastomic homoplasmic lines (pNLyc#2 and LCe) were grown directly on soil. Plants were grown for three weeks in a controlled environment (100–250 μmol m^-2^ s^-1^, 23 °C) and then transferred to fully controlled (plant chamber/530 and 53μmol m^-2^ s^-1^ red and white light respectively, 16/8 h photoperiod, 70 % relative humidity and 24 °C), semi-controlled (greenhouse/average light intensity: 170–380 μmol m^-2^ s^-1^, maximum light intensity: 1200 μmol m^-2^s^-1^ and 24 °C), and uncontrolled conditions (polytunnel/natural climate conditions during spring-summer 2019 in Potsdam, Germany). In each climate condition, plants were grown side by side and they were randomly distributed with at least 50 cm of space between each other. Physiological parameters, such as plant height and leaf and fruit number, were recorded through development (10 to 60–70 days of growth under the different climate conditions) and/or before performing the biomass experiment. Plant biomass for plants grown in fully controlled conditions was assessed in 11-week-old plants (only the biomass of the aerial part, leaf and stem, was recorded). Plant (leaves and stem) and fruit biomass for plants grown under semi-controlled conditions was assessed in two groups of 8- and 16-week-old plants, respectively. The first group was grown for quantification of the leaves and stem (*n*=5-6), and the second was grown for the assessment of fruit biomass (*n*=5). Both groups were grown in parallel and harvested at different time points (eight and 16 weeks, respectively). The biomass of plants grown under uncontrolled conditions in the polytunnel was measured in 12-week-old tomato plants. In this case, the leaf, stem and fruit biomass was recorded from the same plants. Briefly, leaves, stem, and fruit were separated and the fresh weight was recorded immediately. Subsequently, the leaves, stem, and fruit were dried at 70 °C for five days, and the dry weight was recorded. Five (biomass) to ten (plant height) biological replicates were used for each experiment under the different climate conditions. For fruit size quantification, the area of three fully ripened fruit detached from three different greenhouse-grown 16-week-old tomato plants was quantified using ImageJ software.

### Photosynthesis measurements

Wild type and transgenic lines were raised from seeds and grown for three weeks under fully controlled conditions in a phytotron (250 μmol photons m^-2^ s^-1^, 16 h/8 h day/night, 22 °C day/18 °C night, 70% relative humidity; pots of 7 cm diameter). The plants were then transferred to the greenhouse (16 h/8 h day/night regime, 450–800 μmol photons m^-2^ s^-1^ combination of artificial and sun light, 24 °C, 65 % relative humidity), randomly allocated, and acclimated for four weeks before the photosynthetic measurements (49-day-old plants). Photosynthetic parameters, such as CO_2_ assimilation, conductance, and relative electron transport rate (rETR), were measured with a Li-6400XT portable photosynthesis system equipped with a leaf chamber fluorometer (Li-Cor Inc., Lincoln, NE, USA). The measurements were performed during the mornings on fully expanded leaves under growth light conditions (greenhouse, 450 µmol (photons) m^-2^s^-1^ of PAR), with the amount of blue light set at 10% of the photosynthetically active photon flux density to optimize stomatal aperture. The reference CO_2_ concentration was set at 400 µmol CO_2_ mol^-1^ air. All measurements were performed using a 2 cm^2^ leaf chamber maintained with a block temperature of 25°C and a flow rate of 300 mmol air min^-1^. The rETR was calculated according to the method described in (Krall and Edwards, 1992). In addition, total non-photochemical quenching (NPQT), (ΦPSII), (ΦNPQ), and (ΦNO) were measured in the same plants with a MultiSpec (Photosync) instrument (Kuhlgert et al., 2016; Tietz et al., 2017). All measurements were conducted during the early morning (9:00–11:00 am) in the same part of the 7^th^ leaf from seven-week-old plants (growing in 20 cm diameter pots). Five to 12 plants were used for the measurements.

### Water deficit and salinity treatments

Water deficit and salinity treatments were performed under greenhouse conditions. Tomato seeds were sown and raised under control conditions in a phytotron. After three weeks, the seedlings were transferred to the greenhouse and acclimated for four days. The plants were randomized and placed 30 cm apart. For water deficit experiments, control plants (wt and transgenic) were watered once per day (50–200 mL per plant, depending of their water requirements), whereas stressed plants were not watered. Plant height and leaf number were recorded before the stress treatment was initiated (day 0) and again at day 10 of the stress conditions. Phenotypes were recorded by photography at the same time points. For salinity stress, plants were watered with 100 mL of water or 100 mL salt solution (NaCl 200 mM) once per day for seven days. Plant height and leaf number were recorded at day 0 before the onset of the stress treatment and seven days later. At day seven, the stress treatment was discontinued and all plants were watered with 100 mL water for one more week. The plants were photographed again at two weeks after the stress onset (one week of salt treatment and a subsequent week of water only). All tomatoes were grown in 13 cm diameter pots for the stress experiments in the greenhouse. Five to six biological replicates were used for measurements of control and stress-treated plants.

### Photooxidative stress

Leaf discs (1.2 cm diameter) were floated on water at 10 °C and simultaneously exposed for 18 h to strong white light (photon flux density/PFD, 1200 mmol photons m^-2^ s^-1^) produced by an array of light-emitting diodes. The stressed leaf discs were then placed on wet filter paper for measurement of autoluminescence emission after a 2 h dark adaptation, as previously described (Birtic et al., 2011). The emission signal was imaged with a liquid nitrogen-cooled charge-coupled device (CCD) camera (VersArray 1300B, Roper Scientific), with the sensor operating at a temperature of −110 °C. The acquisition time was 20 min, and on-CCD 2 × 2 binning was used, leading to a resolution of 650 × 670 pixels. As previously shown, the imaged signal principally emanates from the slow decomposition of the lipid peroxides that accumulated in the samples during the oxidative stress treatment (Birtic *et al*., 2011).

### Shelf-life experiments

Tomato fruits (*n*=5) were harvested from 16-week-old wild type and transgenic lines and kept for seven weeks at 23°C and a relative humidity ∼20%. The fruit phenotype was recorded 0, 8, 16, 24, 32, 40, and 48 days after detachment from the plant.

### Microscopy analysis

Fully ripened tomato fruits were detached from 12-week-old tomato plants for further microscopy analysis. Lycopene and β-carotene (Lyc+β-car) were observed with a Leica DM6000B/SP5 confocal laser-scanning microscope (Leica Microsystems, Wetzlar, Germany), following a previously published protocol (D’Andrea et al., 2014). The Lyc+β-car signal was visualized using laser excitation of 488 nm and emission between 400 and 550 nm. The total fluorescence of the generated micrographs was quantified using the ROI function in Fiji software, based on collected data from three different tomato fruits from each line.

### HPLC analysis of pigments

Plastid isoprenoids (chlorophylls, carotenoids, and tocopherols) were extracted and quantified as described previously (Emiliani et al., 2018).

### Hormone quantification

Levels of endogenous phytohormones (cytokinins, auxins, jasmonates, abscisates, gibberellins, and salicylic acid) were determined in five biological replicates of freeze-dried tomato leaves and fruit, according to a modified method described previously (Simura et al., 2018). Briefly, samples containing 1 mg DW of biological material were extracted in an aqueous solution of 50% acetonitrile (v/v). A mixture of stable isotope-labeled standards of phytohormones was added to validate the LC-MS/MS method. Crude extracts were loaded onto conditioned Oasis HLB columns (30 mg/1 ml, Waters) and washed with 30% aqueous acetonitrile. Flow-through fractions containing purified analytes were collected and evaporated to dryness in a vacuum evaporator. The chromatographic separation was performed using an Acquity I class system (Waters, Milford, MA, USA) equipped with an Acquity UPLC® CSH C18 RP column (150 × 2.1 mm, 1.7 μm; Waters). The eluted compounds were analyzed using a triple quadrupole mass spectrometer (Xevo™ TQ-XS, Waters) equipped with an electrospray ionization source. Data were processed with Target Lynx V4.2 software, and final concentration levels of phytohormones were calculated by isotope dilution (Rittenberg and Foster, 1940).

### Metabolite profile analysis

The methyl *tert*-butyl ether (MTBE) extraction buffer was prepared and samples extracted as described by Salem et al. (Salem et al., 2016). For metabolites, the chromatograms and mass spectra were evaluated using ChromaTOF 1.0 (Leco, www.leco.com) and TagFinder v.4. (Luedemann et al., 2008) software, respectively. The mass spectra were cross-referenced using the Golm Metabolome database (Kopka et al., 2005). Data are reported following the standards (**Dataset S1 and S2**) suggested by Fernie et al. (Fernie et al., 2011).

### Lipid profile analysis

After MTBE extraction, the lipid-containing fraction was dried, resuspended, and analyzed by LC-MS. Samples were run in negative and positive mode (**Datasets S3 and S4**). The mass spectra were processed with the Refiner MS 7.5 (Genedata) and Xcalibur software.

### Statistical and data analyses

Statistical analysis was performed using GraphPad Prism (version 5.0) or R environment (version 3.5.2 https://www.R-project.org/). Growth and plant productivity were quantified by conducting a set of several experiments. First, growth curves (based on plant height) for all the transgenic lines and their respective wild types were determined for plants grown under fully controlled (plant chamber), semi-controlled (greenhouse), and uncontrolled (polytunnel/ “field” experiment) conditions. Ten plants were used for each environmental condition (*n*=10). The physiological parameters (plant height, leaf number, fruit number) and plant productivity (fresh and dry matter of leaves, stems, and fruit) were quantified on plants grown under fully controlled (*n*=5), semi-controlled (*n*=5-10), and uncontrolled conditions (*n*=5-10). Fruit fresh and dry matter were quantified for the semi-controlled and uncontrolled conditions. Seed yield was quantified in an independent experiment as the total seed production of 12 transgenic and wild type plants for each genotype. Photosynthetic analysis was performed on plants grown under semi-controlled conditions (*n*=5-12). Water deficit and salinity stress experiments were performed on three-week-old tomato plants grown under greenhouse conditions (*n*=5-6). A non-paired two-tailed Student t-test was performed to compare each transgenic line with their respective wild type using GraphPad Prism software. Pigment, metabolite, lipid, and hormone quantifications were performed on five to six tomato plants grown under semi-controlled conditions. Pigments and hormones (*n*=5) were analyzed with the unpaired two-tailed Student t-test to compare each transgenic line with their respective wild type using GraphPad Prism software. For metabolomics (*n*=5), data mining, normalization, clustering, and graphical representation were performed using R Software. For lipid analysis, the output data were normalized to the internal standard and the amount of dry sample used for the analysis (**Datasets S5 and S6**).

For statistical analysis, the MetaboAnalyst webserver was used (Chong et al., 2019; Pang et al., 2020). The data were auto-scaled and normalized. The differences in the distribution of lipid profiles among the transgenic lines were visually explored by principal component analysis (PCA). The supervised partial least squares discriminant analysis (PLS-DA) was used when the separation obtained with PCA was inadequate. Significant differences were determined among the transgenic lines and their respective wild types with the non-parametric Wilcoxon rank-sum test (*n*=5). The patterns of the lipid species that changed across the groups of samples were further investigated by building heatmaps based on the calculated lipid ratios for the transgenic lines and their respective wild types.

## AUTHOR CONTRIBUTIONS

J.C.M: Conceived the project and the experimental design, performed growth, biomass and yield, salt and drought stress, and fruit shelf-life experiments. J.G.V. and J.C.M.: performed photosynthetic experiments with Li-Cor and Multispec, respectively, and performed metabolite extraction and sample preparation (J.C.M.), and data analysis (J.G.V.). J.M. and S.A.: performed apocarotenoid extraction, sample preparation and data analysis. O.N. and I.P.: performed hormonomics analysis. M.R-C.: performed carotenoid extraction and quantification; S.C. and J.C.M.: performed data analysis from lipidomics and lipid extraction, respectively. M.H.: performed high light stress experiments and lipid peroxide quantification. M.K. and J.C.M.: performed microscopy analysis with assistance of J.C.M. JCM wrote the paper with special input from J.G.V., A.R.F., M.R-C., A.S. and all other coauthors.

## ACKNOWLEDGMENTS

We are grateful to Prof. Dr. Lothar Willmitzer for his support and advice. We thank Prof. Dr. Ralph Bock (Max Planck Institute of Molecular Plant physiology, Golm, Germany) and Dr. Caterina D’Ambrosio (Centro Ricerche Metapontum Agrobios, ALSIA, Italy) for kindly providing the transplastomic pNLyc#2 and LCe seeds and the homozygous nuclear High Caro (H.C.) lines, respectively. We thank Dr. Camila Caldana and Anne Michaelis for providing the GC facility and running the GC samples, respectively, and Maria Rosa Rodriguez-Goberna for technical support related with pigment analysis (supported by grant BIO2017-84041-P from the Spanish AEI). In addition, we thank Hana Martínková and Petra Amakorová for their help with phytohormone analyses. The hormonomics work was funded by the Ministry of Education, Youth and Sports of the Czech Republic (European Regional Development Fund-Project “Plants as a tool for sustainable global development” No. CZ.02.1.01/0.0/0.0/16_019/0000827), and the Internal Grant Agency of Palacký University (IGA_PrF_2021_011).

## DECLARATION OF INETERESTS

The authors declare no competing interests.

